# Delineating the transcriptional landscape and clonal diversity of virus-specific CD4^+^ T cells during chronic viral infection

**DOI:** 10.1101/2022.05.12.491625

**Authors:** Ryan Zander, Achia Khatun, Moujtaba Y. Kasmani, Yao Chen, Weiguo Cui

## Abstract

CD4^+^ T cells responding to chronic viral infection often acquire a dysfunctional phenotype that is characterized by a progressive loss in Th1 differentiation and function, as well as an upregulation of multiple co-inhibitory receptors. Conversely, CD4^+^ T cells, and particularly Tfh cells, gradually increase their production of IL-21 during chronic viral infection, which is critical to sustain humoral immunity and also effector CD8^+^ T cell responses. Recent evidence further indicates that a memory-like CD4^+^ T cell population also develops in the face of persistent infection, although how the transcriptional landscape of this subset, along with the Th1 and Tfh cell subsets from chronic infection, differ from their acute infection counterparts remains unclear. Additionally, whether cell-intrinsic factors such as TCR usage influence CD4^+^ T cell fate commitment during chronic infection has not previously been studied. Herein, we perform single-cell RNA sequencing (scRNA-seq) combined with single-cell T cell receptor sequencing (scTCR-seq) on virus-specific CD4^+^ T cells during chronic lymphocytic choriomeningitis virus (LCMV) infection. We identify several transcriptionally distinct states among the Th1, Tfh, and memory-like T cell subsets that form at the peak of chronic infection, including the presence of a previously unrecognized Slamf7^+^ subset with cytolytic features, and show that the relative distribution of these populations differs substantially between acute and persistent LCMV infection. Moreover, while the progeny of most T cell clones displays membership within each of these transcriptionally unique populations, overall supporting a one cell-multiple fate model, a small fraction of clones display a biased cell fate decision, suggesting that TCR usage may impact CD4^+^ T cell development during chronic viral infection. Importantly, a comparative analysis further reveals both subset-specific and core gene expression programs that are differentially regulated between CD4^+^ T cells responding to acute and chronic viral infection. Together, these data may serve as a useful framework and allow for a detailed interrogation into the clonal distribution and transcriptional circuits underlying CD4^+^ T cell differentiation during chronic viral infection.

## Introduction

CD4^+^ T cells play an essential role in coordinating innate and adaptive immunity against multiple pathogens. Upon antigen encounter and exposure to inflammatory cytokines, CD4^+^ T cells display an inherent capacity to adapt diverse functional fates and facilitate immunity through their secretion of soluble meditators and direct cell interactions with other immune cell populations. In response to either an acute or persistent viral infection, naïve antigen-specific CD4^+^ T cells clonally expand and primarily differentiate into either T helper type 1 (Th1) or T follicular helper (Tfh) cells, which support cellular or humoral responses, respectively (Crotty, 2011; Sheikh and Groom, 2021).

Several recent studies have provided insight into how the fate commitment of CD4^+^ T cells is established early after CD4^+^ T cell priming. Notably, the strength of T cell receptor (TCR) and IL-2R signaling strongly influences this bifurcation decision, with increased TCR and IL-2R signaling generally favoring Th1 lineage commitment (Choi *et al*., 2011; Pepper *et al*., 2011; Johnston *et al*., 2012; Snook, Kim and Williams, 2018). Mechanistically, activation of the TCR and IL-2R signaling pathways result in the induction of Blimp-1, a transcription factor (TF) that is critical for Th1 differentiation and that also potently antagonizes Bcl-6, a transcriptional repressor that is required for Tfh differentiation (Johnston *et al*., 2009; Choi *et al*., 2011; Pepper *et al*., 2011). Additionally, early exposure of activated CD4^+^ T cells to IL-12 and type I interferon (IFN) signaling can promote Th1 cell differentiation by inducing expression of T-bet (Hsieh *et al*., 1993; Manetti *et al*., 1993; Mullen *et al*., 2001; Ray *et al*., 2014), which is the lineage-defining TF that is required for Th1 development (Szabo *et al*., 2000). Th1 cells can then travel to sites of infection where they produce pro-inflammatory cytokines such as IL-2, IFN-γ, and TNF-α, which contribute to the activation of CD8^+^ T cells and macrophages (Sheikh and Groom, 2021). Thus, Th1 cells play a critical role in orchestrating immunity against intracellular pathogens. Conversely, dendritic cell (DC) presentation of antigen and secretion of IL-6 facilitates early Tfh differentiation by inducing Bcl-6 expression and the subsequent upregulation of CXCR5 (Crotty, 2011; Eto *et al*., 2011; Choi *et al*., 2013). Pre-Tfh cells then migrate from the T cell zone to the T cell-B cell border where they can interact with antigen-presenting B cells via ICOS-ICOSL and CD40-CD40L pathways, which are required for fully instilling the Tfh program (Han *et al*., 1995; Choi *et al*., 2011; Crotty, 2011). Mature Tfh cells then provide essential help signals to pathogen-specific B cells to facilitate germinal center (GC) reactions, B cell secretion of high-affinity antibodies, and the generation of memory B cells and long-lived plasma cells (Crotty, 2011).

After the initial T cell expansion phase, the pool of virus-specific CD4^+^ T cells begins to decline in both self-resolving and chronic infections, although the nature of the infection plays a major role in shaping the overall magnitude, subset distribution, and functional capacity of responding CD4^+^ T cells. In acute settings, following pathogen clearance, distinct populations of memory CD4^+^ T cells develop that are well equipped to confer long-lasting protection against re-infection. These memory cells include Th1 and Tfh-like subsets (which may represent T effector memory (Tem) cells) (Marshall *et al*., 2011; Pepper *et al*., 2011; Hale *et al*., 2013), tissue resident memory (Trm) cells that provide rapid protection at frontline sites of infection (Iijima and Iwasaki, 2014; Schreiner and King, 2018), and a less differentiated T central memory (Tcm) cell population that displays an increased capacity to proliferate and produce IL-2 upon secondary challenge (Pepper and Jenkins, 2011). While several studies have provided evidence that differentiated effector Th1 and Tfh cells can give rise to long-lived memory cells, previous work further indicates that a memory precursor population develops among early responding CD4^+^ T cells, and that this precursor population also significantly contributes to the establishment of durable and effective CD4^+^ T cell memory following resolution of the infection (Marshall *et al*., 2011; Pepper *et al*., 2011; Ciucci *et al*., 2019). Notably, similar to that of Tcm cells, this precursor subset displays high expression of CCR7, TCF-1, and Bcl2, and as such was referred to as T central memory precursor (Tcmp) cells (Ciucci *et al*., 2019). Importantly, these Tcmp cells display an augmented capacity to survive the contraction phase of acute infection and generate diverse effector cell populations during a secondary response (Marshall *et al*., 2011; Pepper *et al*., 2011). The development and functional fitness of Tcmp cells has recently been identified to be dependent on the TF Thpok (Ciucci *et al*., 2019), although previous work also indicates that Bcl-6, in addition to its well-established role in facilitating Tfh differentiation, is also important for the generation of Tcmp cells (Pepper *et al*., 2011). Thus, CD4^+^ T cell differentiation during acute viral infection is a well-synchronized process that results in appropriately balanced Th1, Tfh, and Tcmp cell formation at the peak of infection, and these subsets can then give rise to highly functional memory populations.

In contrast to acute settings, CD4^+^ T cells responding to persistent infection or cancer often acquire dysfunctional features, including a diminished capacity to produce pro-inflammatory cytokines that coincides with the upregulation of co-inhibitory receptors such as PD-1, Tim-3, and CTLA-4 (Fuller *et al*., 2004; Brooks *et al*., 2005; Crawford *et al*., 2014). Additionally, several studies have identified that Th1 formation and function is drastically reduced during chronic viral infection, and this decline in Th1 formation is accompanied by a relative increase in Tfh cell development (Fahey *et al*., 2011; Yamada *et al*., 2016). This shift in CD4^+^ T cell differentiation is thought to be important to limit Th1-mediated immunopathology (Penaloza-MacMaster *et al*., 2015), while at the same time promoting Tfh-mediated antibody responses that are required for viral clearance from the periphery (Fahey *et al*., 2011; Harker *et al*., 2011; Cook *et al*., 2015; Greczmiel *et al*., 2017). Notably, our recent work has further demonstrated that in addition to the anticipated Th1 and Tfh populations that form during chronic viral infection, a previously unrecognized memory-like CD4^+^ T cell subset also develops and bears a similar transcriptional program to that of Tcmp cells from acute viral infection (Zander *et al*., 2022). Thus, CD4^+^ T cells responding to chronic viral infection are also capable of exhibiting memory-like properties, even in the presence of ongoing viral replication. We have further demonstrated that although this memory-like subset shows an augmented capacity to accumulate, and does display some developmental plasticity, it largely stays in a quiescent-like state in the face of persistent infection (Zander *et al*., 2022), which may hinder its capacity to provide “help” signals to other immune cell subsets. However, despite these recent advances, our understanding of how CD4^+^ T cells adapt under settings of persistent antigen exposure and inflammation remains incompletely understood. Moreover, whether additional heterogeneous populations of CD4^+^ T cells emerge during chronic viral infection and how these subsets compare to those from acute infection have remained significant knowledge gaps in the field. Additionally, questions remain about how chronic viral infection perturbs the functional and transcriptional landscape of CD4^+^ T cell populations, and whether these alterations in the CD4^+^ T cell compartment are due to global differences in gene expression programs or stem from differences in the distribution pattern of T helper cell subsets that develop under these contexts.

While previous studies have demonstrated that most antigen-specific CD4^+^ T cell clones can give rise to multiple lineages following acute bacterial or viral infection (Tubo *et al*., 2013), recent evidence indicates that some clones may also display a biased cell fate decision (Khatun *et al*., 2021). Importantly, cell-intrinsic factors, such as TCR signal strength, TCR affinity for cognate peptide, and TCR chain usage have all recently been implicated in contributing to shaping CD4^+^ T cell fate decision during viral infection (Tubo *et al*., 2013, 2016; Cho *et al*., 2017; Künzli *et al*., 2020; Khatun *et al*., 2021). Interestingly, TCR signal strength was found to exert opposite effects on the balance of Th1 to Tfh cells depending on whether the infection was acutely resolving or persistent (Künzli *et al*., 2020). However, we currently do not fully understand the extent by which TCR repertoire impacts cell fate decisions during chronic viral infection. Additionally, whether naïve T cells with particular TCR chains display the propensity to give rise to multiple independent lineages, or acquire a particular differentiation program that may skew the T helper cell response in the setting of chronic viral infection, remains unclear.

In this study, we employed single-cell RNA sequencing (scRNA-seq) coupled with single-cell TCR sequencing (scTCR-seq) to investigate the transcriptional landscape of CD4^+^ T cells responding to chronic LCMV infection. Notably, we uncovered previously unappreciated transcriptional heterogeneity among the virus-specific CD4^+^ T cell pool, including the identification of several transcriptionally distinct states within the Th1, Tfh, and T memory-like cell populations. Moreover, we found that, similar to findings from acute viral infection, the majority of GP66-specific CD4 T cell clones responding to chronic viral infection displayed remarkable developmental plasticity as evidenced by their membership across multiple distinct subsets, although a small fraction of clonotypes did appear to exhibit a preferred lineage choice. Lastly, using an integrated analysis to compare the transcriptional program and subset distribution of CD4^+^ T cells responding to either acute or chronic LCMV infection, we identified that persistent infection is not only associated with altered CD4^+^ T cell subset formation and gene expression profiles, but also changes in core gene expression modules that span multiple distinct T helper cell populations.

## Results

### GP66-specific CD4^+^ T cells responding to chronic LCMV infection are transcriptionally diverse and are dominated by a few large clones

To better understand the transcriptional heterogeneity and clonal diversity among CD4 T cells responding to chronic viral infection, we performed paired single-cell RNA sequencing (scRNA-seq) and single-cell T cell receptor sequencing (scTCR-seq) on splenic GP_66-77_ (GP_66_)_-_specific CD4 T cells that were isolated from two individual mice on day 10 post-LCMV Clone 13 (Cl13) infection. We defined a clone as a group of T cells sharing the same nucleotide sequence from the CDR3 regions of both the α and β chains of the T cell receptor (TCR). In order to safeguard against potential errors produced during FACS sorting or sequencing misreads, we restricted our analysis to clones with at least two cells sharing the same TCR α and β chain CDR3 sequences. This resulted in a total of 168 clones for mouse 1 (M1) and 159 clones for M2 (**Figure 1A**), consisting of 5,185 total cells between our two samples (2836 cells for M1, 2349 cells for M2). These findings are in line with the expected range of TCR clonotypes reported in mice (Kotturi *et al*., 2008; Jenkins and Moon, 2012; Khatun *et al*., 2021) and suggest that our scRNA-seq approach likely recovered the majority of the constituents from the GP_66_-specific clonal repertoire. Interestingly, and similar to our analysis on the T cell repertoire for GP_66_-specific CD4^+^ T cells during acute LCMV Armstrong infection, we did not observe any clonal overlap between M1 and M2 when α and β chain CDR3 sequences were paired on a per cell basis, although a few instances of TCR CDR3 overlap was observed when considering either α and β chain CDR3 sequences individually (**Figure 1B**). These results highlight a tremendous amount of diversity within the TCR repertoire, even among genetically identical hosts. We next assessed patterns of clonal distribution and dominance. Notably, the top five clones for each mouse (which ranged from comprising 3.7% to 9.8% of the GP_66_-specific pool) accounted for ~40-45% of all cells in the dataset (**Figure 1A, C**), demonstrating that the CD4^+^ T cell response during LCMV Cl13 infection is dominated by a few large clones.

**Figure 1:**
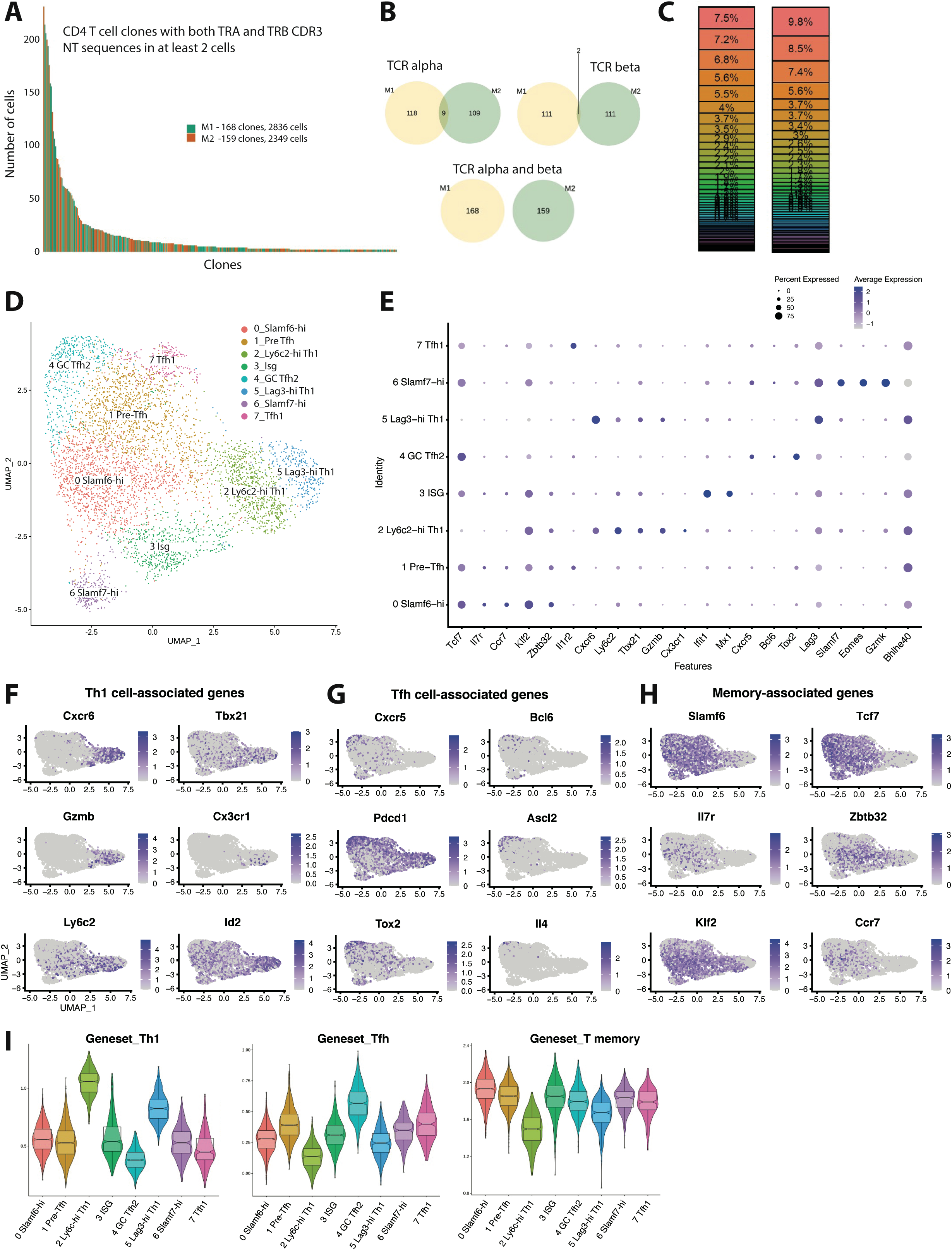
Clonal distribution and transcriptional landscape of GP66-specific CD4+ T cells during chronic LCMV infection. **(A-I)** GP66-specific splenic CD4 T cells were isolated and sort-purified from two individual LCMV Cl13-infected mice on day 10 p.i. and the 10X genomics pipeline was used to generate a combined scRNA-seq + TCR-seq library. A downstream computational analysis was then performed in R. (A). Summary data showing the size of and clonal distribution of GP66-specific CD4+ T cells (with at least 2 cells per clone) recovered from the two separate mice. (B). Pie charts showing the shared number of public clones when assessing the sequence of either TCR alpha chains, TCR beta chains, or both TCR alpha and beta chains. (C). Summary data showing the relative frequency of each T cell clone recovered from the two mice. (D). UMAP plot showing the presence of 8 transcriptionally distinct clusters of GP66-specific CD4 T cells. (E). Dot plot showing the relative expression of differentially expressed genes between the 8 clusters. (F-H). UMAP plots depicting the expression of either Th1 cell-associated genes (F), Tfh cell-associated genes (G), or memory-associated genes (H). (I). Module score analyses were performed to compare the relative Th1, Tfh, and memory T cell signatures among the distinct clusters of GP66-specific CD4+ T cells.

Next, using uniform manifold approximation and projection (UMAP) to visualize single-cell gene expression data, we identified the presence of eight transcriptionally distinct clusters stemming from our dataset containing ≥ 2 cells per clone of GP_66_-reactive CD4^+^ T cells (**Figure 1D, S1A**). Of these eight clusters, clusters 2 and 5 both displayed increased expression of several Th1 cell-associated genes, such as *Cxcr6, Tbx21* (encodes T-bet), *GzmB*, and *Id2* (Shaw *et al*., 2016; Choi *et al*., 2020) (**Figure 1E-F**), suggesting that these two populations are of the Th1 lineage. Interestingly, cluster 2 displayed increased expression of *Ly6c2, Klf2, S1pr*1, and also displayed uniquely high expression of *Cx3cr1*, whereas cluster 5 displayed increased expression of *Lag3, Slamf1, Pdcd1*, and *Tnf* (**Figure 1E, S1B**). These findings may suggest the presence of functional heterogeneity within the Th1 pool, and we henceforth refer to these populations as the Ly6c-hi Th1 and Lag3-hi Th1 clusters, respectively. Using Seurat’s module score feature to more accurately quantify the gene expression patterns of these clusters in comparison to published gene sets further supported that cluster 2 and cluster 5 are comprised of Th1 cells (**Figure 1I**). By contrast, clusters 4 and 7 displayed the lowest Th1 module scores, and also displayed increased expression of Tfh-associated genes such as *Cxcr5* and *Bcl6* (**Figure 1E, G, I**). Cluster 4 in particular displayed high expression of genes important for GC Tfh differentiation and function, including *Ascl2, Tox2*, and *Il4* (Crotty, 2014; Liu *et al*., 2014; Xu *et al*., 2019) (**Figure 1G**). Consistent with these observations, cluster 4 displayed the highest Tfh module score (**Figure 1I**), followed by cluster 7, the latter of which also displayed a transcriptional profile similar to the T-bet–expressing Tfh subset that develops during acute LCMV Armstrong infection (Khatun *et al*., 2021) (**Figure S1C**). Thus, we referred to clusters 4 and 7 as GC Tfh2 and Tfh1 cell subsets, respectively.

Clusters 0 and 1 both exhibited high expression of *Slamf6* (encodes Ly108), as well as multiple memory-associated genes, including *Tcf7, Il7r, Ccr7*, and *Klf2* (Ciucci *et al*., 2019) (**Figure 1E, H**). Consistent with this observation, Clusters 0 and 1 also displayed increased memory T cell module scores (**Figure 1I**), and conversely lower Th1 and Tfh module scores, although Cluster 1 did display the third highest Tfh module score behind the GC Tfh2 and Tfh1 clusters. Together, these data suggest that cells from clusters 0 and 1 likely fall within the *Slamf6*^+^ T memory-like cell subset that we recently demonstrated develops during LCMV Cl13 infection and largely remains in a quiescent-like state, yet does display some developmental plasticity, including the potential to give rise to Tfh cells in the midst of persistent infection (Zander *et al*., 2022). Interestingly, compared to Cluster 0, Cluster 1 did display increased expression of *Tnfrsf4, Batf, Il1r2*, and *Cd83* (**Figure 1E, S1A**), which were also highly expressed in the Tfh1 population, possibly indicating that cells from cluster 1 are in a more activated state. Given that Cluster 1 had an increased Tfh module score relative to Cluster 0, coupled with our finding that some memory-like CD4^+^ T cells can give rise to Tfh cells (Zander *et al*., 2022), we elected to refer to Clusters 0 and 1 as Slamf6-hi and Pre-Tfh cells respectively.

Lastly, our scRNA-seq analysis identified the presence of two additional transcriptionally distinct clusters, cluster 3 and cluster 7, with cluster 3 cells displaying uniformly high expression of multiple type I IFN stimulated genes, including *Ifit1, Ifit3, Isg15, Isg20, Mx1*, and *Cxcl10* (**Figure 1E, S1A**). On the other hand, cluster 7 cells displayed uniquely high expression of *Slamf7, Eomes*, and *Gzmk* (**Figure 1E, S1A**), and as such, cells within this cluster bore a resemblance to a cytolytic CD4^+^ T cell subset that has been identified in tumors as well as autoimmune disease (Della-Torre *et al*., 2018; Cachot *et al*., 2021). Notably, and consistent with their diverse transcriptional programs, the eight clusters we identified also substantially differed in their relative expression of several cytokines and chemokines (**Figure S1D**). Although *Ifng* was expressed by all clusters, Ly6c-hi Th1 and Lag3-hi Th1 clusters displayed the highest expression of *Ifng*, and these respective clusters also had increased expression of the Th1-associated chemokine ligands (Ccl) *Ccl3, Ccl4*, and *Ccl5* (**Figure S1D**). Conversely, and consistent with our previous report (Zander *et al*., 2022), *Il21* expression was increased in the pre-Tfh, Tfh1, and GC Tfh2 populations (**Figure S1D**), whereas *Cxcl10*, a chemokine known to be induced by type I IFN signaling (Groom and Luster, 2011), was enriched in the ISG cluster. Intriguingly, the Slamf7-hi subset displayed increased expression of the immunoregulatory chemokine *Tgfb1*, although all eight clusters expressed *Tgfb1* to some extent. Taken together, these data uncover broad transcriptional and functional heterogeneity among CD4 T cells responding to chronic LCMV infection.

### Validation of CD4^+^ T cell heterogeneity during chronic LCMV infection

Using several differentially expressed surface markers identified in our scRNA-seq analysis, we next performed spectral flow cytometry to further investigate the phenotypic heterogeneity within the CD4 T cell compartment during LCMV Cl13 infection. Using a panel of 18 parameters, t-distributed stochastic neighbor embedding (t-SNE) visualization was performed on PD-1^hi^ CD4^+^ T cells (which include both GP_66_-specific and other LCMV epitope-reactive CD4^+^ T cells) that were obtained from LCMV Cl13-infected Mx1-GFP reporter mice. Notably, several surface molecules were differentially expressed amongst the pool of LCMV-specific CD4^+^ T cells, which allowed for the formulation of several distinct clusters (**Figure 2A**). Consistent with our scRNA-seq analyses, a subset of cells displayed low expression of Ly108 and high expression of CXCR6, indicating that these cells are likely of the Th1 lineage (**Figure 2A**). Additionally, one small population (which only comprised ~ 2-6% of the CD4 T cell pool), displayed uniformly and exclusively high expression of Slamf7 (**Figure 2A**), indicating that these cells likely correspond with the *Slamf7*-hi cells from cluster 6 in our scRNA-seq analysis. Another subset displayed increased expression amounts of CXCR5 and PD-1 relative to the other cells, suggesting that this population is likely comprised of cells found within the GC Tfh2 subset. Interestingly, some cells that clustered together exhibited intermediate expression amounts of CXCR5 and high expression of IL-1R2 (**Figure 2A**), a marker that was increased in both the Pre-Tfh and Tfh1 subsets, possibly indicating that IL-1R2 expression can be used to distinguish these populations of CD4^+^ T cells. A large proportion (~36%) of cells expressed high amounts of Ly108 and IL-7Ra, suggesting that this population of cells likely corresponds with the *Slamf6*-hi cells from cluster 0. Of note, cells did not appear to cluster together based off of Mx1-GFP expression; rather, Mx1, an interferon-stimulated protein, was detected via GFP expression across all t-SNE clusters (**Figure 2A**). Given that type I IFN-stimulated genes remain elevated for several weeks following chronic viral infection (Ref), this finding suggests that all subsets of CD4^+^ T cells are likely to experience some degree of type I IFN signaling during persistent viral infection. In contrast to our findings at the protein level, we likely detected a discrete cluster of IFN-stimulated CD4^+^ T cells by scRNA-seq because IFN-stimulated genes transiently dominated the transcriptomes of certain cells, causing them to cluster together.

**Figure 2:**
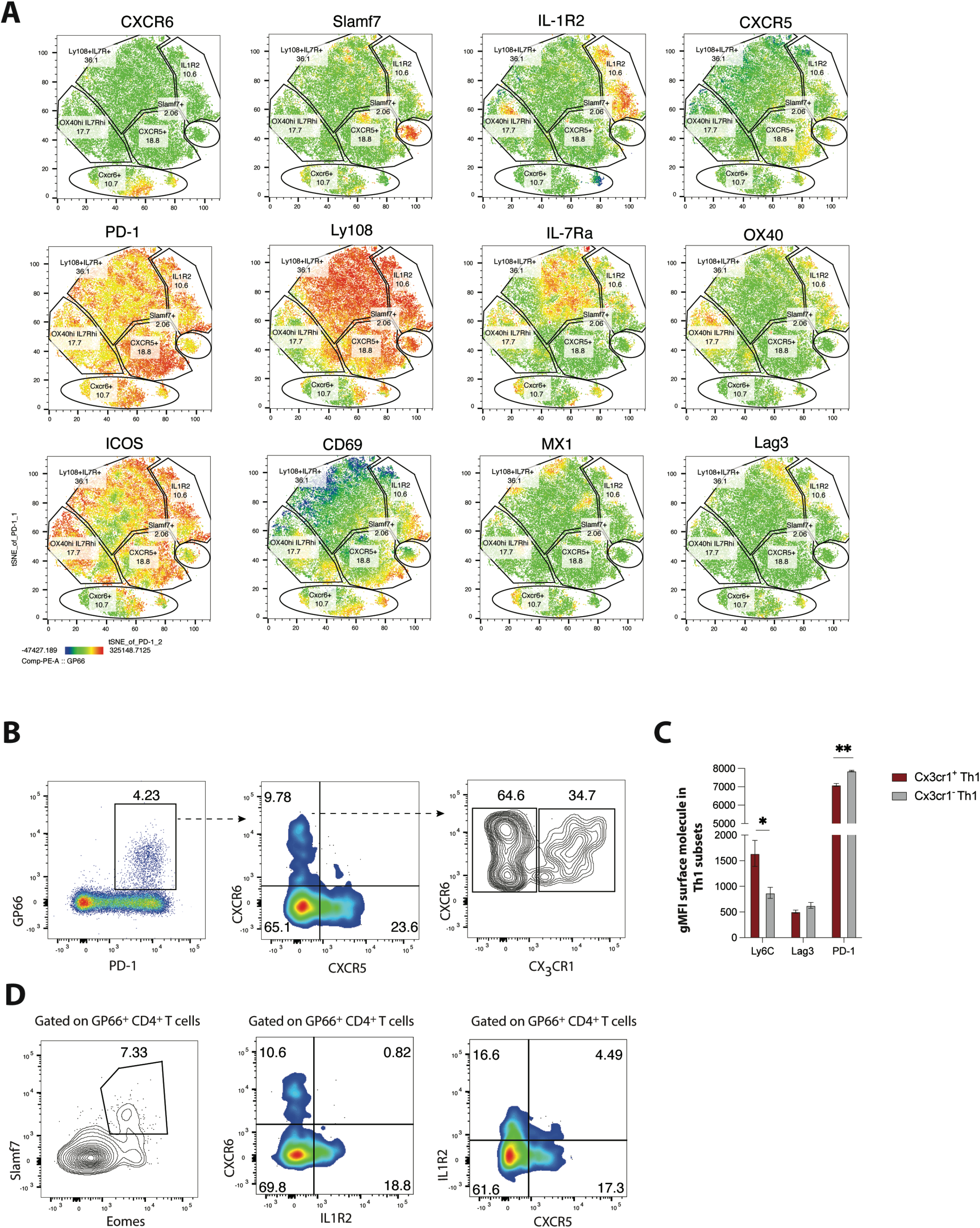
Validation of heterogeneity in CD4 T cells responding to chronic LCMV infection using Spectral flow cytometry. (A). MX1-GFP reporter mice were infected with LCMV Cl13 and spectral flow cytometry was used to assess the phenotype of splenic CD4 T cells on day 25 p.i. Using a panel of 18 parameters, t-distributed stochastic neighbor embedding (t-SNE) visualization was performed on total PD-1^hi^ CD4 T cells (which include both GP66-specific and other LCMV epitope-reactive CD4 T cells). Representative tsne flow plots in A are displayed for one mouse, with data being representative of 3 individual mice. (B). Representative flow plots showing potential gating strategy to identify CX3CR1+/− CXCR6+ Th1 cell subsets. (C) Summary data showing the relative expression of indicated surface molecules in CX3CR1+ and CX3CR1− Th1 cell subsets. (D). Representative flow plots showing potential gating strategies to identify Slamf7+Eomes+ (far left), and IL-1R2+CD4 T cell subsets (which may be comprised of preTfh and Tfh1 subsets). Data (mean ± S.E.M.) in Figure 2C are from 3 individual mice, and are representative of at least two independent experiments.

Recently, we demonstrated that differential expression of CXCR6 and CXCR5 can be used to distinguish three major CD4^+^ T cell subsets that differ in their phenotype, function, transcriptional program and epigenetic landscape: CXCR6^hi^ CXCR5^lo^ CD4 T cells exhibit a robust Th1 cell program, CXCR6^lo^ CXCR5^hi^ CD4 T cells correspond with Tfh cells, and CXCR6^lo^ CXCR5^lo^ cells (which have high Ly108, IL-7R, and TCF-1 expression) display a memory-like signature and as such were defined as a T memory-like cell subset (Ref). Given the additional heterogeneity uncovered in this study, we next aimed to determine some potential gating strategies that can be used to further distinguish some of the major CD4^+^ T cell subsets identified in our scRNA-seq analyses. Notably, and consistent with our scRNA-seq findings, GP_66_-specific CXCR6^+^ Th1 cells could be further stratified based on CX_3_CR1 expression, with CX_3_CR1^hi^ CXCR6^hi^ cells displaying increased expression of Ly6C and lower expression of PD-1 compared to CX_3_CR1^lo^ CXCR6^hi^ cells (**Figure 2B-C**). Given that distinct Th1 subsets can emerge during different contexts and have diverse functional roles (Jenkins), a more in-depth investigation into the functional heterogeneity of these Th1-like populations may have important implications for diseases where Th1 cells play a dominant role.

Using a different gating strategy, we detected a minor population of GP_66_-specific CD4^+^ T cells that displayed high co-expression amounts of Eomes and Slamf7 (**Figure 2D**), with these markers being largely unique to the *Slamf7*^hi^ subset identified in our scRNA-seq analysis. Thus, high expression of Slamf7 may be a reliable surface marker for this particular subset of CD4^+^ T cells. Similarly, a clear population of IL-1R2-expressing CD4^+^ T cells was detected amongst the GP_66_-specific pool (**Figure 2D**), with the majority of these cells staining negative for CXCR6 and also exhibiting low CXCR5 expression, possibly suggesting that this expression module may potentially mark the pre-Tfh subset. Interestingly, a small fraction (~4-5%) of GP_66_-specific CD4 T cells coordinately expressed both IL-1R2 and CXCR5, possibly marking cells from the Tfh1 cluster (**Figure 2D**), which our scRNA-seq analyses identified as expressing both of these molecules (**Figure 1E, G**). Collectively, our results highlight several potential gating strategies and markers that may be useful in demarcating the populations identified in our scRNA-seq analysis, which may allow for further characterization of these transcriptionally and phenotypically distinct subsets.

### Trajectory analysis at the clonal level demonstrate developmental plasticity among most GP_66_-specific CD4^+^ T cell clones during chronic LCMV infection, although some clones do display a biased-cell fate decision

To gain insight into the potential developmental relationships between these cell subsets, we employed use of the monocle package in R (Qiu *et al*., 2017). Monocle aligns cells based on gene expression levels to determine cell trajectories in psuedotime, a measure of how much progress an individual cell has made along a differentiation pathway (Qiu *et al*., 2017). Our data formed a tripartite tree under Monocle analysis, with cells from the Ly6c-hi Th1 and Lag3-hi Th1 clusters residing on the far left branch, cells from the GC Tfh2, Tfh1, and Slamf7 clusters dwelling on the far right branch, and cells from the Slamf6-hi subset localizing more on the central branch (**Figure 3A**). Notably, cells from both the Slamf6-hi and Pre-Tfh clusters could readily be detected along all three branches (**Figure 3A-B**), possibly indicating that these subsets may retain a higher degree of developmental plasticity. Interestingly, whereas ~ 19.3% of Slamf6-hi cells were found to reside on the Tfh branch, ~55.6% of cells from the pre-Tfh cluster were found on this branch (**Figure 3A-B**), further supporting that cells from the pre-Tfh cluster may be en route towards developing along a Tfh differentiation pathway.

**Figure 3:**
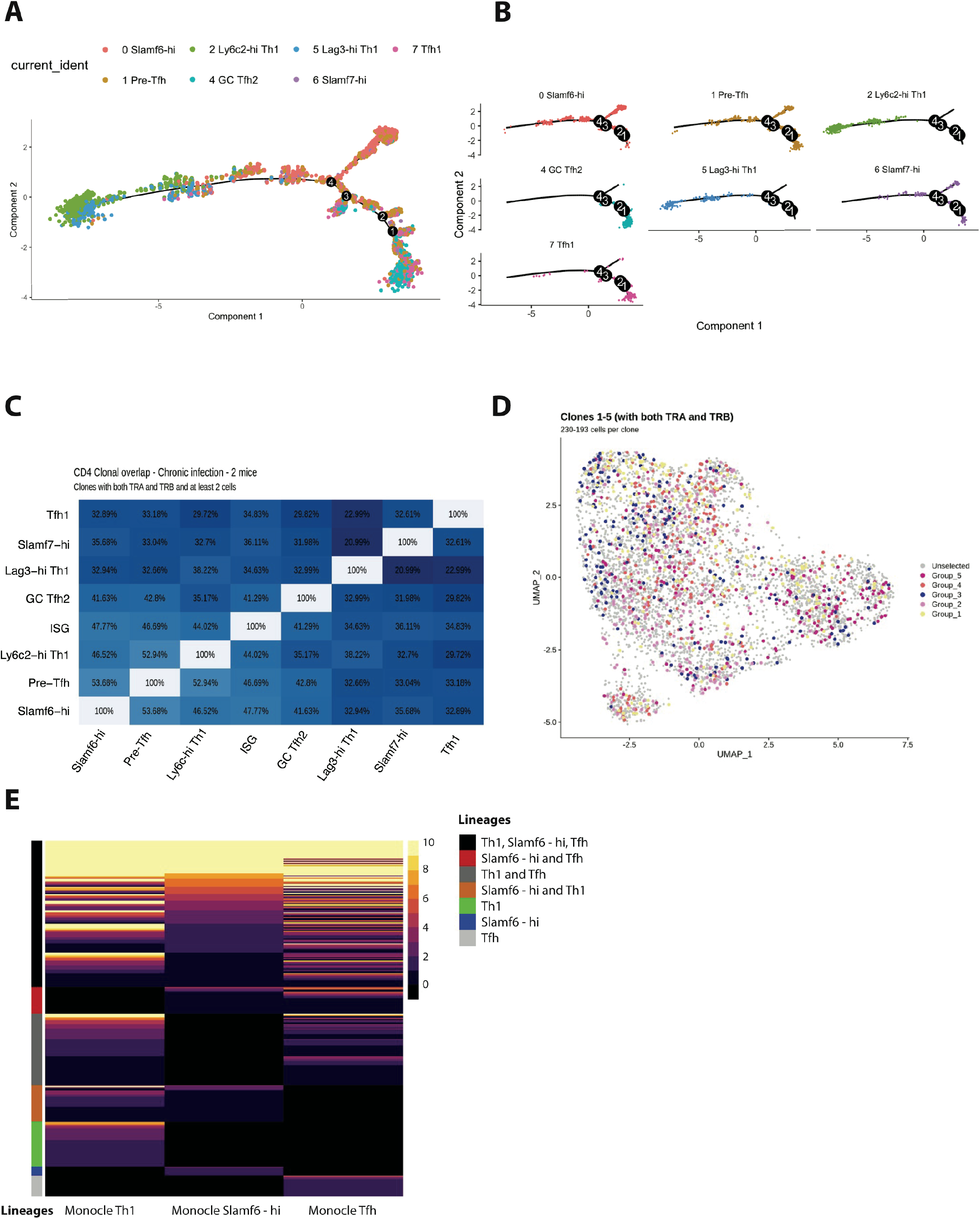
Trajectory analysis and clonal lineage tracing highlights that most GP66-specific CD4 T cells display developmental plasticity during chronic LCMV infection, although a small fraction of clones exhibit a biased cell fate decision. (A-B). Monocle tree trajectory plots showing predicted cellular differentiation of GP66-specifc CD4 T cell subsets based on trajectory analysis. (C). Summary chart displaying clonal overlap among GP66+ CD4 T cell subsets. (D). UMAP plot as in Figure 1D, but colored by clone membership. (E). Heatmap showing subset distribution and frequency of 329 clones (n=168 clones from M1 and n=159 clones from M2) among indicated CD4 T cell lineage (Th1, Tfh, and Slamf6^hi^ memory) as defined by Monocle state.

Previous work indicates that in response to acute viral or bacterial infection, the majority of pathogen-specific CD4^+^ T cell clones can give rise to both Th1 and Tfh cells following clonal expansion (Tubo *et al*., 2013; Becattini *et al*., 2015; Cho *et al*., 2017). Accordingly, we recently reported that a sizeable fraction (~28%) of CD4^+^ T cell clones display a preferred lineage choice toward either Th1 or Tfh cells during acute LCMV Armstrong infection (Khatun *et al*., 2021). Notably, TCR structure, and in particular the CDR3 motif of the TCR α chain, was determined to be a driving influencer of this biased cell fate decision (Khatun *et al*., 2021). However, given that TCR signaling strength may exert opposite effects on the balance of Th1 to Tfh cells depending on the nature of the infection (Künzli *et al*., 2020), whether CD4^+^ T cells responding to chronic viral infection also display similar patterns of differentiation at the clonal level remains unclear. Merging our single-cell clonal and transcriptomic datasets together allowed us to track the lineage commitment of CD4^+^ T cell clones from the GP_66_-specific repertoire during chronic LCMV Cl13 infection. Notably, a high degree of clonal overlap was observed amongst all CD4^+^ T cell subsets identified in our scRNA-seq analysis, with most subsets displaying >30% overlap with any given subset (**Figure 3C**). In line with this finding, upon examination of the top 5 clones (which contained 193-230 cells per clone) we identified that each clone displayed membership in every cluster from our scRNA-seq analysis (**Figure 3D**), suggesting that similar to acute infection, one naïve CD4^+^ T cell displays the capacity to give rise to multiple T helper cell lineages during persistent infection. Intriguingly, when assessing the cellular distribution of all clones with at least 2 cells per clone (168 clones for M1 and 159 clones for M2), we identified that while the majority of clones (~75%) had some membership in at least two or more lineages (Th1, Slamf6-hi, or Tfh as defined by Monocle branch), a small fraction of clonotypes exhibited a preferred lineage choice, with ~ 15% of clones displaying select membership in only the Th1 lineage and < 10% of clones displaying exclusive membership in either the Slamf6-hi or Tfh lineages (**Figure 3E**). Thus, similar to our observations in acute LCMV infection (Khatun *et al*., 2021), while the majority of GP_66_-specific CD4^+^ T cell clones appear to display some developmental plasticity, a minor proportion of clones do appear to display a biased cell-fate decision.

### CD4^+^ T cells responding to chronic LCMV infection display altered subset distribution patterns and core changes in their transcriptional program

Global (bulk) transcriptional analysis previously performed on total GP_66_-specific CD4^+^ T cells isolated from mice either acutely or persistently infected with LCMV identified distinct patterns of gene expression, with CD4^+^ T cells responding to chronic infection displaying increased expression of genes encoding PD-1, Lag3, and CTLA-4 co-inhibitory receptors, as well as increased expression of several transcription factors, including *Eomes, Prdm1* (encodes Blimp-1), *Zbtb32*, and *Klf4* (Crawford *et al*., 2014). However, as multiple transcriptionally distinct subsets have since been found to develop during acute and chronic viral infection, whether these observed alterations in the transcriptional program of CD4^+^ T cells stem from differences in the relative distribution of CD4^+^ T cell subsets that develop under different contexts, or from a conserved gene expression program acquired by CD4^+^ T cells responding to chronic viral infection remains unclear. To gain insight into this question, we performed a comparative scRNA-seq transcriptomic analysis by combining our previously published scRNA-seq data of GP_66_^+^ CD4^+^ T cells isolated from LCMV Armstrong-infected mice on day 10 p.i. (Khatun *et al*., 2021) with our current dataset of GP_66_^+^ CD4 T cells from day 10 post-LCMV Cl13 infection. To do this, we used the integration function in Seurat to perform a canonical correlation analysis (CCA) (Butler *et al*., 2018) and identify shared subpopulations across datasets. Notably, all of the clusters initially identified in our LCMV Cl13-restricted dataset (**Figure 1**) were also detected in this integrated dataset, although clear differences in the relative proportion of each cluster was observed when grouped by infection status (Acute vs Chronic; **Figure 4A**). In particular, the proportion of Ly6c2-hi Th1 and Lag3-hi Th1 cell subsets were dramatically reduced (by ~50%) during LCMV Cl13 as compared to LCMV Armstrong infection; this reduction in Th1 cell formation was accompanied by increased frequencies of Slamf6-hi, Pre-Tfh, ISG, and Slamf7-hi clusters, whereas GC Tfh2 and Tfh1 clusters remained similar between infections (**Figure 4A-B**).

**Figure 4:**
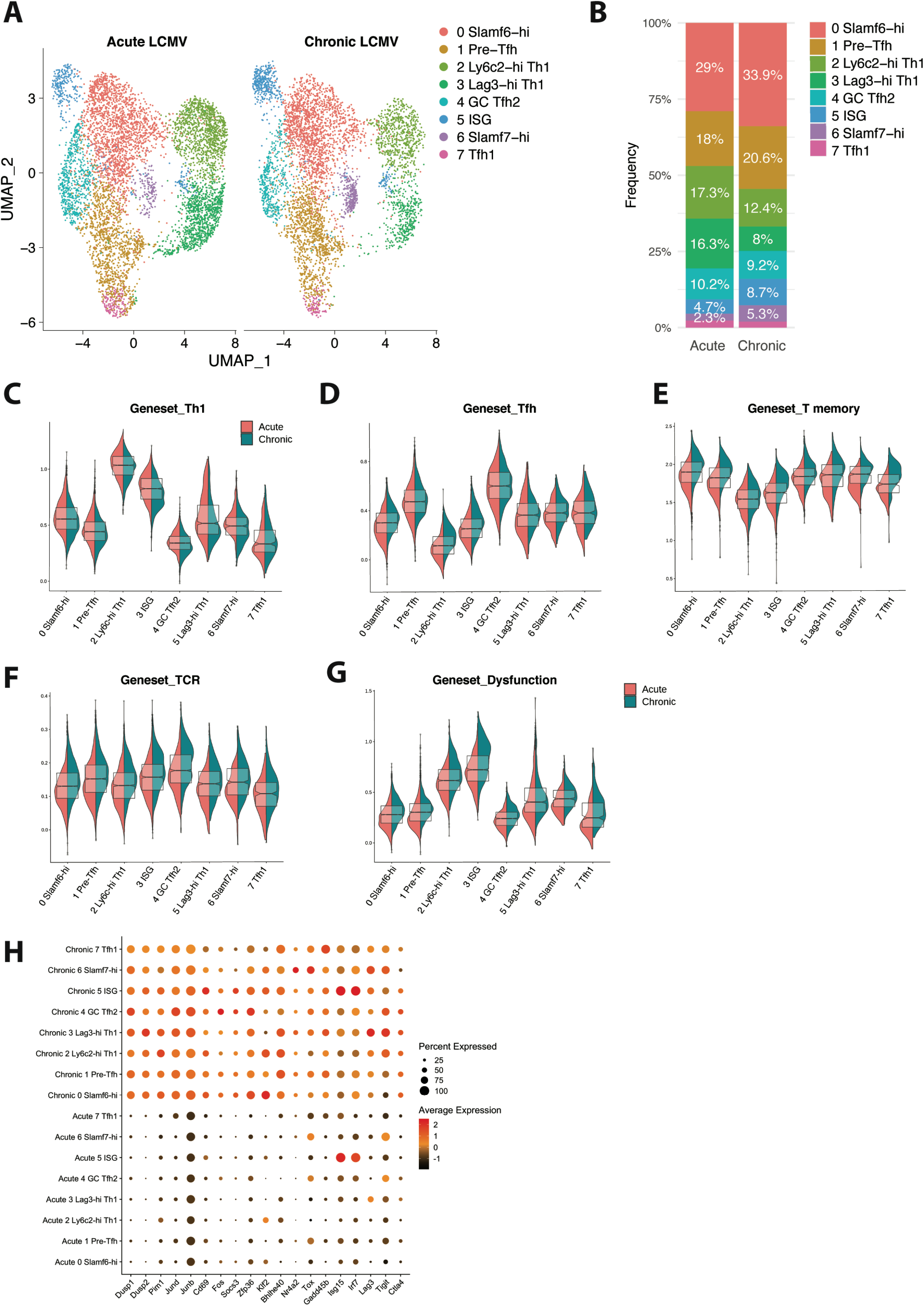
CD4 T cells responding to chronic viral infection display altered subset distribution patterns and core gene expression modules compared to their acute infection counterparts. (A). A canonical correlation analysis (CCA) was performed on scRNA-seq libraries obtained from GP66-specific CD4+ T cells during chronic LCMV Cl13 infection (this study) and acute LCMV Armstrong infection (Khatun et al.) that were integrated together using the integration function in Seurat (Butler et al. 2018). (B). Summary graph showing the relative subset distribution of GP66+ CD4 T cells subsets during acute and chronic LCMV infection on day 10 p.i.. (C-G) Module score analyses showing the average expression levels of the Th1, Tfh, memory T cell, TCR signaling, and T cell dysfunction programs among the CD4 T cell subsets identified in A. (H). Summary dot plot showing the average gene expression and the proportion of each cell subset expressing indicated gene. All genes listed were among the top 50 differentially expressed genes that were upregulated across multiple distinct subsets in response to chronic LCMV infection.

This observation is in line with previous work demonstrating a rapid loss of Th1 cell formation and function during chronic viral infection (Fahey *et al*., 2011; Yamada *et al*., 2016), as well as our recent finding that this decline in CXCR6^+^ Th1 cell accumulation is accompanied by an increase in the formation of a Slamf6^+^ memory-like CD4^+^ T cell subset (Zander *et al*., 2022). In addition, our finding that the ISG cluster is overrepresented during LCMV Cl13 infection is consistent with previous reports showing that type I IFN-stimulated genes remain elevated during chronic viral infection compared to acute infection (Teijaro *et al*., 2013; Wilson *et al*., 2013). Interestingly, although Eomes was previously identified to be overexpressed in CD4^+^ T cells during chronic infection (Crawford *et al*., 2014), in this study we identified that this is likely due to a preferential expansion of the Slamf7^+^ subset, of which displays uniquely high expression of Eomes (**Figure 1E, 2D**). Consistent with this observation, we found that the proportion of Slamf7^+^ Eomes^+^ virus-specific CD4^+^ T cells was increased by ~10-fold during chronic LCMV Cl13 infection compared to acute LCMV infection (**Figure S2A**). Taken together, these data indicate that the nature of the infection (i.e. resolution vs chronicity) may impact the relative distribution of CD4^+^ T cell subsets that develop, with persistent infection resulting in a notable attenuation in Th1 responses and an accompanying increase in the formation of memory-like, pre-Tfh, ISG, and Slamf7^+^ CD4^+^ T cell populations.

To assess potential shifts in gene expression profiles that varied by infection and that spanned across all CD4^+^ T cell subsets, we performed module score analyses to compare Th1, Tfh, memory T cell, TCR signaling, and T cell dysfunction gene signatures across all of the CD4^+^ T cell clusters. Notably, module score analysis demonstrated that most CD4^+^ T cell clusters from chronic LCMV infection exhibited decreased expression of the Th1-associated program, possibly suggesting that an attenuation in Th1-like activity is a broad feature of CD4^+^ T cells responding to chronic viral infection and not only a result of a decrease in the Th1 lineage (**Figure 4C**). Conversely, Tfh module scores were generally similar between acute and chronic LCMV CD4^+^ T cell clusters, although the Ly6c2-hi Th1 and Lag3-hi Th1 cell subsets from chronic LCMV infection did display a relatively increased Tfh signature compared to their acute infection counterparts (**Figure 4D**). Intriguingly, although the relative score varied by cluster, both memory T cell and Tcmp gene signatures were enriched across all CD4^+^ T cell clusters stemming from chronic LCMV infection (**Figures 4E, S2B**), possibly suggesting that persistent infection promotes a core quiescence-or memory-associated transcriptional program in responding CD4^+^ T cells, which may be important to limit Th1-mediated immunopathology (Penaloza-MacMaster *et al*., 2015). This observation is in line with our previous finding that CXCR6^+^ Th1, CXCR5^+^ Tfh, and memory-like CD4^+^ T cell subsets all display equivalent accessibility at key memory-associated gene loci, such as *Il7r, Tcf7, Slamf6*, and *Ccr7*, indicating that multiple distinct CD4^+^ T cell lineages may remain poised to upregulate a memory-associated program during chronic infection (Zander *et al*., 2022). Additionally, and similar to CD8^+^ T cells responding to chronic infection (Wherry *et al*., 2007), all CD4^+^ T cell clusters from LCMV Cl13 infection uniformly displayed increased TCR signaling and T cell dysfunction gene expression profiles (**Figure 4F-G**), once again suggesting that persistent exposure to antigen and inflammation may result in the upregulation of some conserved gene expression programs that span multiple phenotypically distinct CD4^+^ T cell subsets. In line with this, all CD4^+^ T cell clusters from LCMV Cl13 infection displayed notable increases in their expression of certain intracellular signaling phosphatases and kinases, such as *Dusp1, Dups2*, and *Pim1*, as well in their expression of several genes encoding various TFs, including *Jund, JunB, Fos, Klf2*, and *Bhlhe40* (**Figure 4H**). Additionally, genes known to be downstream of TCR signaling such as *Cd69, Nr4a2* and *Tox* (Alfei *et al*., 2019; Chen *et al*., 2019; Khan *et al*., 2019), interferon-stimulated genes such as *Isg15* and *Irf7*, and genes encoding the inhibitory receptors Lag3, Tigit, and CTLA4 were all overexpressed across all CD4^+^ T cell clusters from chronic infection compared to acute LCMV infection (**Figure 4H**). Together these data indicate that certain core transcriptional programs are upregulated across virus-specific CD4^+^ T cell subsets independent of phenotype in response to sustained exposure to persistent viral infection. Conversely, some TFs with increased expression during chronic infection are more uniquely expressed by particular CD4^+^ T cell subsets, such as *Zbtb32* in the Slamf6-hi cluster, *Eomes* in the Slamf7-hi cluster, and *Prdm1* (encodes Blimp1; **Figure S2C**) in the Th1 clusters from chronic infection, indicating that some gene expression programs are specific to a particular lineage. Of note, several genes were also found to be downregulated in particular CD4^+^ T cell subsets during chronic infection, such as *Id2* and *Cxcr6* in Th1 cell clusters as well as *Cxcr5* and *Icos* in the GC Tfh2 cluster, whereas Id3 appeared to be downregulated in most CD4^+^ T cell subsets from chronic LCMV infection (**Figure S2C**). Collectively, these results demonstrate that chronic viral infection results in both altered CD4^+^ T cell subset distribution as well as distinct gene expression programs, with CD4^+^ T cell subsets displaying not only lineage-specific gene expression profiles but also core gene expression programs that are conserved across multiple distinct populations of T helper cells.

## Discussion

In this study, we demonstrate that CD4^+^ T cells responding to chronic LCMV infection are more heterogeneous than previously appreciated, with several transcriptionally distinct subsets being detected within Th1, Tfh, and T memory-like CD4^+^ T cell populations. We further demonstrate that while the progeny of most CD4^+^ T cell clones displayed immense developmental plasticity, suggesting a one cell-multiple fate model, a small fraction of clones appeared to exhibit a biased cell fate-decision, indicating that TCR usage may have an impact on CD4^+^ T cell differentiation. Additionally, we highlight some potential flow cytometric gating strategies that can be used to further interrogate these respective subsets and also provide evidence that the relative distribution of these populations differed substantially between acute and chronic viral infection. Lastly, our integrated analyses revealed that chronic viral infection results in vastly altered gene expression programs in responding CD4^+^ T cells, with evidence for not only lineage-specific gene expression profiles but also core gene expression programs that are upregulated and conserved across multiple distinct populations of T helper cells. Collectively, these data may serve as a useful resource to assess and compare the transcriptional landscape and differentiation trajectory of virus-specific CD4^+^ T cell clones during acute and chronic viral infection.

Several recent studies have identified that CD4^+^ T cells responding to acute viral infection are comprised of multiple transcriptionally distinct CD4^+^ T cell subsets, including Th1, Tfh, and Tcmp cells (Ciucci *et al*., 2019; Khatun *et al*., 2021). While the differentiation and helper functions of Th1 and Tfh subsets in these acute settings is fairly well-established, recent work has also yielded important insight into the transcriptional profile and developmental biology of Tcmp cells. Notably, Tcmp cells display high expression of memory cell markers CCR7, TCF-1, and Bcl-2 and have an intrinsic capacity to survive the contraction phase of acute infection and generate diverse effector cell populations during a secondary response (Marshall *et al*., 2011; Pepper *et al*., 2011; Ciucci *et al*., 2019). The development of Tcmp cells has been demonstrated to be dependent on both Thpok and Bcl6 (Pepper *et al*., 2011; Ciucci *et al*., 2019). Thus, our understanding of CD4^+^ T cell differentiation during acute infection is becoming increasingly more well-defined. By comparison, our knowledge of how CD4^+^ T cells adapt in the face of persistent antigen exposure and inflammation remains incompletely understood. Moreover, whether diverse populations of CD4^+^ T cells also emerge during chronic infection, or whether they can display memory-like properties have remained significant knowledge gaps in the field. Notably, our recent work has identified that a memory-like CD4^+^ T cell population develops alongside Th1 and Tfh cell subsets during chronic viral infection, and that this memory-like subset bears a strikingly similar transcriptional profile as Tcmp cells from acute LCMV infection (Zander *et al*., 2022). However, whether additional heterogeneity exists amongst the CD4^+^ T cell pool during chronic viral infection remains unclear. Moreover, how the transcriptional landscape of CD4^+^ T cells differs between acute and chronic viral infection at the single-cell level has not previously been explored. Herein, we demonstrate that CD4 T cells responding to chronic viral infection are more heterogeneous than previously appreciated, with several transcriptionally distinct subsets arising during the early phase of chronic infection. These include two potentially unique Th1 cell subsets (Ly6c-hi and Lag3-hi cells), two Tfh-like subsets (GC Tfh2 and Tfh1), a memory-like population, and a pre-Tfh subset that exhibits both a memory-like signature and also expresses some Tfh-associated genes. Additionally, our scRNA-seq analyses identified a cluster of Slamf7-hi cells that may represent a cytolytic CD4 T cell subset (Della-Torre *et al*., 2018; Cachot *et al*., 2021), as well as a unique cluster that displayed increased expression of multiple ISGs, consistent with chronic infection driving a sustained type I IFN response (Teijaro *et al*., 2013; Wilson *et al*., 2013). Using spectral flow cytometry, we validate the formation of several of these subsets during chronic viral infection, and also highlight a few potential gating strategies and markers that may be useful in interrogating these subsets further. However, future work will be important to elucidate the developmental relationship between these respective populations and to establish whether some of these newly identified subsets represent functionally distinct lineages or merely transitional states along the Th1, Tfh, or memory T cell developmental pathways.

Notably, while our TCR clonal trajectory analyses indicate that the majority of GP66-specific CD4^+^ T cell clones responding to chronic LCMV infection appear to be multipotent, a small fraction of T cell clones appeared to display a biased cell fate decision and favored a particular lineage. Overall, this finding is consistent with other recent studies from resolving infections (Tubo *et al*., 2013; Khatun *et al*., 2021), and suggests that the vast majority of naïve CD4^+^ T cells are capable of giving rise to multiple distinct lineages independent of TCR sequence, although TCR structure may still influence the differentiation trajectory of some select clones. As certain CDR3 motifs from the TCR alpha chain have previously been implicated as being a useful predictor of T cell fate during acute LCMV infection (Khatun *et al*., 2021), a deeper investigation into whether particular TCR alpha or beta chain sequences impact CD4^+^ T cell fate determination during chronic viral infection will be of future interest.

Consistent with chronic viral infection resulting in diminished Th1 differentiation and function (Fuller *et al*., 2004; Brooks *et al*., 2005; Fahey *et al*., 2011; Yamada *et al*., 2016), our integrated scRNA-seq analysis demonstrated that the proportions of both Ly6c-hi and Lag3-hi Th1 cells were dramatically reduced during chronic versus acute LCMV infection. Interestingly, module score analyses identified that loss of the Th1 gene expression program was not restricted to only the Th1 cell subsets but was conserved across multiple distinct subsets, indicating that loss of Th1-like pro-inflammatory activity may be a generalizable feature of T helper cell responses during chronic viral infection. This loss in Th1 cell subset formation coincided with increased frequencies of multiple subsets, including Slamf6-hi, Pre-Tfh, ISG, and Slamf7-hi clusters, whereas the Tfh clusters remained similar between infections. Intriguingly, we further observed an enriched quiescence or memory-associated transcriptional profile across most CD4^+^ T cell subsets responding to chronic viral infection, despite viral load remaining high at this time point. Although the reasons for this remain unclear, it is possible that CD4^+^ T cells responding to chronic infection need to acquire a more quiescent or memory-like phenotype in order to mitigate Th1-mediated pathology (Penaloza-MacMaster *et al*., 2015). This enhanced acquisition of a memory-like phenotype is supported by the relative increased formation of Slamf6^+^ and pre-Tfh populations during persistent infection, which exhibit high expression of *Tcf7, Il7r, Ccr7*, and *Bcl2*. However, it is important to note that despite this enriched memory-associated program, most CD4^+^ T cell subsets responding to chronic LCMV infection also displayed increased TCR signaling and T cell dysfunction module scores, indicating that these cell subsets are not likely to be in a truly quiescent state, nor do they fit the description of canonical memory populations. Consistent with this, we identified that *Id3*, a TF known to be critical for memory T cell formation following acute infection (Yang *et al*., 2011), was substantially diminished in most CD4^+^ T cell subsets from chronic infection compared to their acute infection counterparts. Moreover, in line with increased TCR signaling during chronic viral infection, we observed a conserved increase in the expression of AP-1 family members *Jund* and *Junb*, as well as in *Cd69, Nr4a2*, and *Tox* gene expression across all CD4^+^ T cell subsets during LCMV Cl13 infection. Similarly, exhaustion markers *Lag3, Tigit*, and *Ctla4* were uniformly increased in CD4^+^ T cell subsets responding to chronic viral infection, consistent with these populations displaying increased T cell dysfunction scores. Collectively, our results highlight perturbed gene expression programs in CD4^+^ T cells during chronic viral infection, with apparent alterations in both lineage-specific T helper cell programs and also core expression modules that are differentially regulated across all CD4^+^ T cell subsets between acute and chronic viral infection. A future investigation into the transcriptional circuits underlying the altered CD4^+^ T cell differentiation that occurs during chronic infection may have important implications for strategies aimed at improving T cell-based immunotherapies during chronic infection.

## Methods

### Mice and LCMV Cl13 infection

Six-to eight-week-old female C57Bl/6 mice obtained through the National Cancer Institute grantees program (Frederick, MD) were used for all experiments, unless otherwise indicated. Mx1-GFP homozygous mice (Jackson strain #:033219) were crossed to C57Bl/6 mice to generate Mx1-GFP heterozygous mice that were then used for some experiments where indicated. Mice were bred and maintained in a closed breeding facility, and mouse handling conformed to the requirements of the Institutional Animal Care and Use Committee guidelines of the Medical College of Wisconsin. Mice were infected with 2×10^6^ PFU of lymphocytic choriomeningitis virus (LCMV) strain Clone 13 (Cl13) via retroorbital injection to establish chronic viral infection.

### Flow cytometry and cell sorting

For cell sorting experiments, splenic CD4^+^ T cells from LCMV Cl13-infected mice were isolated using the EasySep mouse CD4 T cell isolation kit (STEMCELL; Cat#19852). Enriched CD4^+^ T cells were then stained using LCMV-specific GP_66-77_ PE tetramer from National Institutes of Health) along with CD4 and CD44 antibodies in FACS buffer. The staining was performed in the dark for 1 hour at room temperature, followed by three washes with FACS buffer. Gp66-specific CD4^+^ T cells were sort-purified on a FACS-Aria sorter. For FACS analyses, single cell suspensions obtained from spleenocytes from LCMV Cl13-infected mice were stained with GP_66-77_ PE tetramer in conjunction with other surface antibodies in FACS buffer, followed by three washes in FACS buffer. Samples were then acquired on either an Aurora Cytek (Cytek Biosciences) or LSR-II (BD Biosciences). In some experiments, after surface staining, cells were fixed with buffer from the True-Nuclear Transcription Factor Buffer Set (BioLegend) for one hour. Then cells were then washed with permeabilization buffer and stained with antibodies against transcription factors in permeabilization buffer. Analyses were performed using Flowjo software version 10.8.1.

| **Flow Cytometry Reagents** |  |  |
| --- | --- | --- |
| APC/Fire anti-mouse CD4 (GK1.5) | Biolegend | Cat #100460<br>RRID:AB_2572111 |
| AF647 anti-mouse CD4 (GK1.5) | Biolegend | Cat #100460<br>RRID:AB_2572111 |
| Pe/Cyanine7 anti-mouse PD.1 (RMP1-30) | Biolegend | Cat #109110<br>RRID:AB_572017 |
| APC/Cyanine7 anti-mouse PD.1 (29F.1A12) | Biolegend | Cat #135224<br>RRID:AB_2563523 |
| APC/Cyanine7 anti-mouse/human CD44 (IM-7) | Biolegend | Cat #103028<br>RRID:AB_830785 |
| APC anti-mouse CXCR5 (L138D7) | Biolegend | Cat #145506<br>RRID:AB_2561970 |
| PE/Dazzle 594 anti-mouse CXCR6 (SA051D1) | Biolegend | Cat #151117<br>RRID:AB_2721700 |
| PerCP-eFluor710 anti-mouse CXCR6 (SA051D1) | Biolegend | Cat #RRID:AB_ |
| BV785 anti-mouse Lag3 | Biolegend | Cat #125219<br>RRID:AB_256657 |
| PE/Cyanine7 anti-mouse CD127 (A7R34) | Biolegend | Cat #135014<br>RRID:AB_1937265 |
| Brilliant Blue 700 Tim3 | BD Biosciences | Cat #747619<br>RRID:AB_2744185 |
| Spark NIR 685 anti-mouse CD69 | Biolegend | Cat #104557<br>RRID:AB_2860607 |
| BUV395 anti-mouse CD44 | BD Biosciences | Cat #740215<br>RRID:AB_2739963 |
| BV711 anti-mouse PSGL1 | BD Biosciences | Cat #740746<br>RRID:AB_2740414 |
| BV650 anti-mouse CD83 | BD Biosciences | Cat #740619<br>RRID:AB_2740317 |
| BV605 anti-mouse PD-1 | Biolegend | Cat #135220<br>RRID:AB_2562616 |
| BV510 anti-mouse Ly108 | BD Biosciences | Cat #745073<br>RRID:AB_2742691 |
| PerCP cy5.5 anti-mouse CX3CR1 (SA011F11) | Biolegend | Cat #149010<br>RRID:AB_2564494 |
| PerCP anti-mouse Ly-6c | Biolegend | Cat #128028<br>RRID:AB_10897805 |
| Pacific Blue anti-mouse Bst2 | Biolegend | Cat #127108<br>RRID:AB_2028455 |
| BV421 anti-mouse Slamf7 | BD Biosciences | Cat #747997<br>RRID:AB_2872458 |
| Anti-mouse IL1R2 | Thermofisher | Cat #PA5-47759<br>RRID:AB_11152697 |
| Pecy5 anti-mouse ICOS | Biolegend | Cat #107708<br>RRID:AB_313337 |
| APC-Fire750 anti-mouse OX40 |  | Cat #119423<br>RRID:AB_2715993 |
| PE LCMV I-A(b) GP66-77 tetramer | NIH Tetramer Core Facility | https://tetramer.yerkes.emory.edu |

### scRNA-seq

LCMV-specific (GP_66-77_ tetramer^+^) CD4^+^ T cells were harvested from the spleens of two LCMV Cl13–infected mice on day 10 after infection and were FACS sorted using a FACS-Aria Sorter (BD Biosciences). Sorted cells were loaded onto the 10X Chromium Controller with a target cell number of 5,000 per mouse. scRNA-seq libraries were prepared using the Chromium Single Cell 5′ v2 Reagent Kit (10X Genomics) according to the manufacturer’s protocol. Two libraries were then quantified using the Kapa Library Quantification Kit and then were loaded onto an Illumina NextSeq 500 sequencer with the NextSeq 500/550 High Output Kit v2.5 (150 cycles; 20024907; Illumina) with the following conditions: 26 cycles for read 1, 98 cycles for read 2, and 8 cycles for the i7 index read. Raw sequencing data were downloaded from Illumina BaseSpace, then demultiplexed and converted to gene-barcode matrices using the “mkfastq” and “count” functions in Cell Ranger v3.0 (10x Genomics). A total of 4394, and 3613 cells were recovered from samples M1 and M2 respectively.

### scTCR-seq

A 10 μL aliquot of each sample’s amplified cDNA was taken for scTCR-seq. The standard 10x protocol was followed for TCR gene amplification. Libraries were quantified using a KAPA library quantification kit (Roche Sequencing) and then loaded onto an Illumina NextSeq 500 sequencer using a NextSeq 500/550 High Output Kit v2.5 (150 cycle kit) (20024907, Illumina) with 150 cycles for read 1, 150 cycles for read 2, and 8 cycles for i7 index reads. Raw sequencing data were downloaded from Illumina BaseSpace, then demultiplexed and converted to gene-barcode matrices using the “mkfastq” and “vdj” functions in Cell Ranger v3.0 (10x Genomics).

### Combined analysis of scRNA-seq and scTCR-seq data

Analysis was primarily performed in R (v 3.6.1) using the package Seurat (v 3.1) (Butler et al., 2018; Stuart et al., 2019), with the package tidyverse (v 1.2.1) (Wickham et al., 2019) used to organize data and the package ggplot2 (v 3.2.1) used to generate figures. scRNA-seq data sets were integrated and then scTCR-seq data was added. scRNA-seq data was filtered to keep cells with a low percentage of mitochondrial genes in the transcriptome (< 5%) and between 200 and 3000 unique genes to exclude poor quality reads and doublets. Cell cycle scores were regressed when scaling gene expression values and T cell receptor genes were regressed during the clustering process, which was performed with the Louvain algorithm within Seurat and visualized with UMAP. Cells other than CD4^+^ T cells, sample-specific outliers, and cells not belonging to clones with at least 2 cells were excluded. A clone was defined as a group of cells sharing the same TCR α and β chain CDR3 nucleotide sequences. Trajectory analyses were performed using monocle (v 2.12.0) (Qiu et al., 2017). The Monocle tree structure was generated using the DDRTree algorithm with the top 100 differentially expressed genes from each Seurat cluster.

### Gene set enrichment and module score analyses

A preranked analysis module in GSEA (Subramanian et al., 2005) was used and the gene sets were found in MSigDB (Liberzon et al., 2011). For single cluster enrichment analysis, differentially expressed genes were identified first (logFC >0.1) and the average logFC expression of each gene was calculated. Then, this average gene expression data for each cluster was used as an input for GSEA analysis. After this, a gene set variation analysis score (Hänzelmann et al., 2013) was calculated using log normalized expression data for each cluster and chosen gene sets, as identified by GSEA analysis. Seurat clusters were scored by GSEA gene sets using the Addmodulescore function. Gene sets for Th1, GC T_FH_, Tcmp, T cell dysfunction, and memory CD4 or CD8 T cells were obtained from published data (Ciucci et al., 2019, and Wherry et al. 2007). Module scores for TCR signaling were calculated within our Seurat clusters using the “T cell receptor signaling pathway” gene set from GSEA (KEGG hsa04660).

### Cellular distribution at the clonal level for three lineages

The cellular distribution of all clones (two or more cells/clone; *n* = 327 clones) among the three different lineages defined by Monocle states (Th1, Tfh, and Slamf6 memory-like; described above) was shown using the pheatmap R package (Fig. 3 E). This allowed for visualization of cellular distributions of each clone among the lineages, thus. demonstrating the differentiation preference of each clone (i.e., single fate or multifate).

### Canonical Correlation Analysis

A comparative scRNA-seq transcriptomic analysis was performed by combining our previously published scRNA-seq data of GP_66_^+^ CD4^+^ T cells isolated from two LCMV Armstrong-infected mice on day 10 p.i. (Khatun et al. 2020 GSE 158896) with our current dataset of GP_66_^+^ CD4 T cells from day 10 post-LCMV Cl13 infection. To do this, we used the integration function in Seurat to perform a canonical correlation analysis (CCA) (Butler et al. 2018) and identify shared subpopulations across datasets. As an initial step of the analysis, cells were filtered based on having high percentage of mitochondrial genes in the transcriptome (> 10%) and less than 200 to ~ 4,000-5,000 unique genes to remove any doublets and dead cells. Following this, a total of ~ 18,000 cells were included in the analysis for four mice. Cell cycle gene scoring was calculated for all cells and regressed out. To understand cellular heterogeneity, unsupervised clustering was performed using a dimension reduction algorithm called, Uniform manifold approximation and projection (UMAP), based on 3000 highly variable genes with an input of the top 20 principle components.

## Data availability

The scRNA-seq and scTCR-seq data have been deposited in the GEO database (accession no GSE201730), and are available to the public.

## Statistical analysis

Statistical tests for flow cytometry data were performed using Graphpad Prism 7. p-values were calculated using two-tailed unpaired Student’s t-tests. P values for violin plots showing gene set–specific module scores across clusters were calculated by the Wilcoxon test with Holm-Sidak correction.

## Acknowledgements

This work is supported by NIH grants AI125741 (W.C.), AI148403 (W.C.), DK127526 (M.Y.K), AI153537 (R.Z.), and by an American Cancer Society (ACS) Research Scholar Grant (W.C.); and by an Advancing a Healthier Wisconsin Endowment (AHW) Grant (W.C). R.Z. also received support from a Cancer Research Institute Irvington Fellowship. M.Y.K. is a member of the Medical Scientist Training Program at the Medical College of Wisconsin (MCW), which is partially supported by a training grant from NIGMS (T32-GM080202). This research was completed in part with computational resources and technical support provided by the Research Computing Center at MCW.

## Author Contributions

R.Z., A.K. and W.C. conceptualized and designed the study. R.Z., A.K., M.Y.K., and Y.C., performed experiments and performed data analyses. R.Z. wrote the manuscript. A.K., M.Y.K., and W.C. edited the manuscript. W.C. and R.Z. supervised the study.

## Figure Legends

**Supplemental Figure 1:**
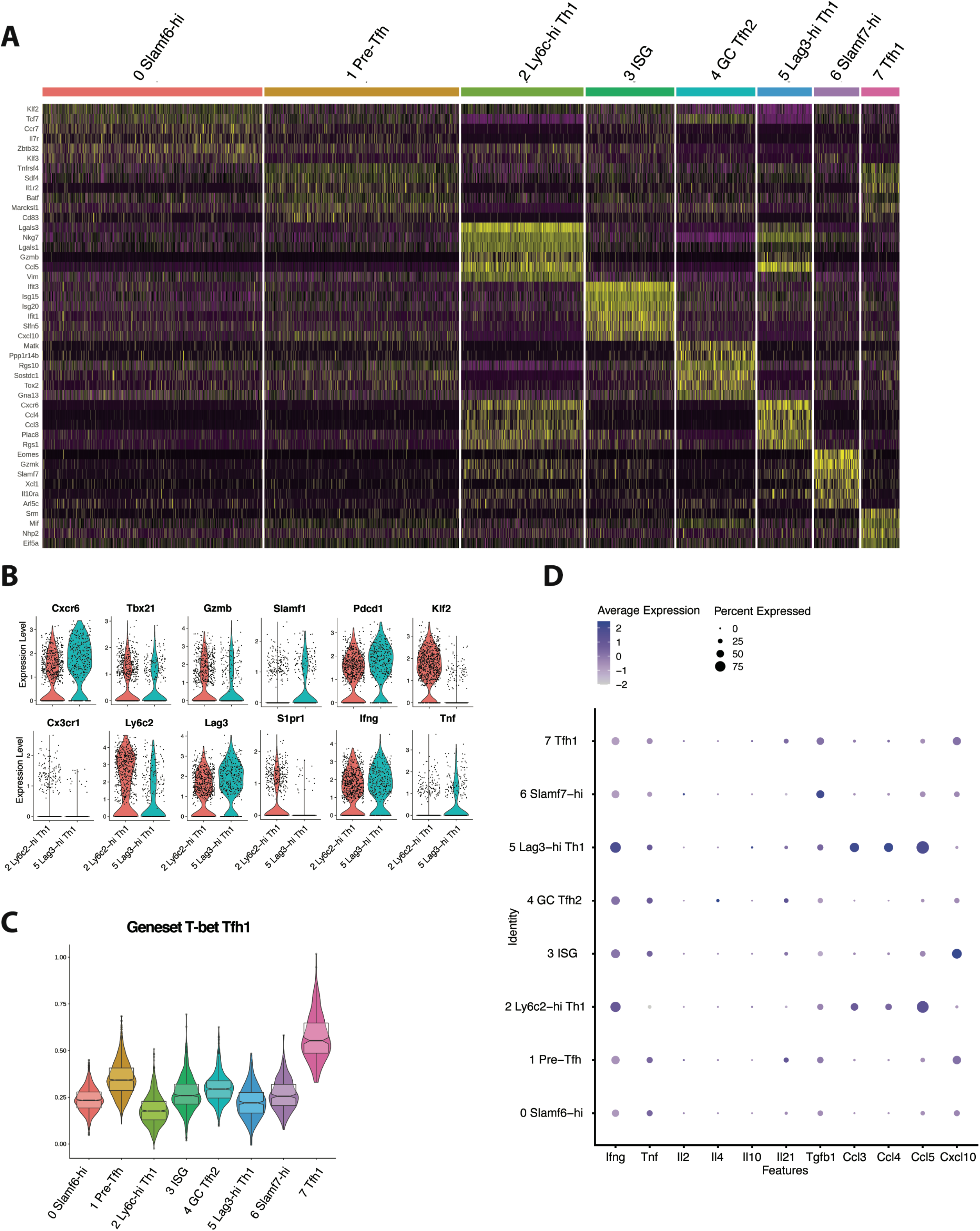
CD4^+^ T cells responding to chronic viral infection are comprised of several transcriptionally and functionally distinct populations. (A). Heatmap showing the top differentially expressed genes in all eight clusters. (B). Violin plots displaying the relative expression of indicated genes in Ly6c-hi and Lag3-hi Th1 cell subsets. (C). Module score analysis showing the average expression levels of the previously identified T-bet Tfh1 program in the eight clusters identified in Figure 1D. (D). Dot plot summary showing the average expression of indicated cytokines and chemokines, and the percent of CD4^+^ T cell subsets expressing each molecule.

**Supplemental Figure 2:**
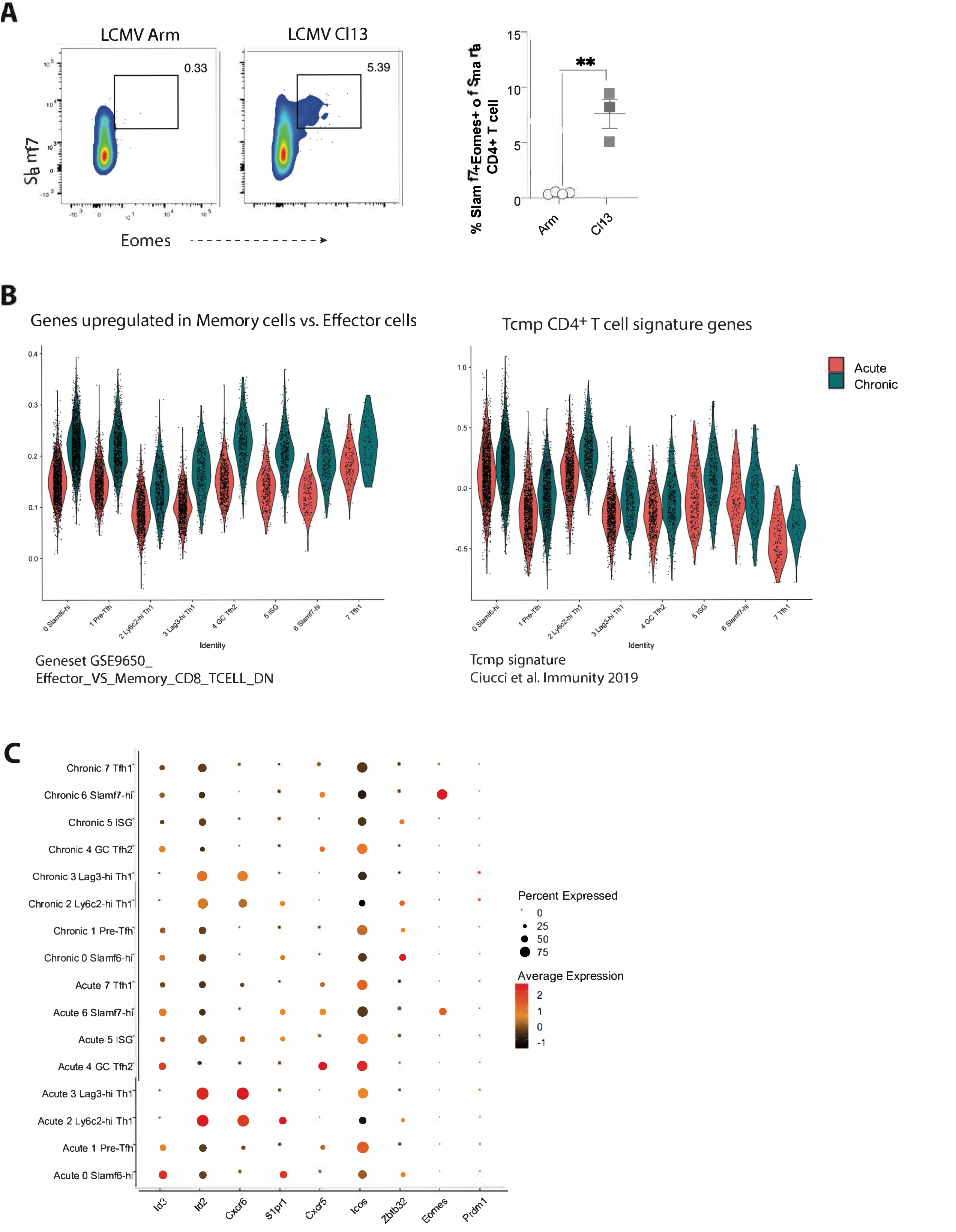
CD4 T cells responding to chronic LCMV infection display altered subset distribution patterns and changes in gene expression programs compared to their acute infection counterparts. (A). Representative flow plots (left) and summary data (right) showing the proportion of the Slamf7^+^Eomes^+^ subset among TCR Tg Smarta CD4+ T cells on day 14 p.i. (B). Module score analyses showing the average expression levels of genes upregulated in memory cells vs. effector cells (left), or the Tcmp signature (right) in each of the 8 clusters identified in our scRNA-seq analysis. (C). Summary dot plot showing the relative expression of each indicated gene and the proportion of each CD4+ T cell subset expressing the respective gene. Data (mean ± S.E.M.) in Figure S2A are from 3 mice, and are representative of at least two independent experiments.

